# State-dependent photocrosslinking unveiled the role of intersubunit interfaces on Ca^2+^ activating mechanisms of the BK channel

**DOI:** 10.1101/2022.12.15.520511

**Authors:** Alberto J. Gonzalez-Hernandez, Belinda Rivero-Pérez, David Bartolomé-Martín, Diego Álvarez de la Rosa, Andrew J.R. Plested, Teresa Giraldez

## Abstract

BK channels are high-conductance potassium channels that are activated by voltage and Ca^2+^. The pore-forming α-subunits form homotetramers including a membrane-spanning domain and a cytosolic domain with a tandem of RCK-like domains (RCK1 and RCK2) per subunit. The eight RCKs compose high affinity Ca2+-binding sites that drive channel activation. Full-length Cryo-EM structures revealed intersubunit interactions between the RCK domains. Asparagine 449 (N449, human BK channel) is located at the RCK1 domain and coordinates Ca^2+^ in the RCK2 of the adjacent subunit. In addition, two Arginines (R786 and R790) of one RCK2 and Glutamate 955 of the adjacent subunit constitute an additional interaction interface. Functional studies on these residues showed that these two interfaces are crucial in Ca^2+^ sensitivity. To detect structural rearrangements induced by Ca^2+^ during channel activation, we took advantage of the photoactivatable unnatural amino acid p-benzoyl-L-phenylalanine (BzF). Functional channels were obtained with this amino acid inserted at 11 positions. N449 and R786 positions (N449BzF and R786BzF respectively). UV-induced photocrosslinking led to Ca^2+^ dependent and voltage-independent effects in both mutants. N449BzF showed a steady-state current reduction at saturating Ca^2+^ concentrations. Our data shows that this effect mainly relies on full occupancy of the RCK1 Ca^2+^ binding site, since mutation of this site abolished the effect. The R786BzF construct showed a substantial potentiation of the current in the absence of Ca^2+^. In this case, photocrosslinking seems to favor the activation of the channel by voltage. Overall, these results suggest mobile interfaces between RCK domains are key to BK channel activation.

## Introduction

The large conductance voltage- and Ca^2+^-activated K^+^ channel (BK or slo1) is constituted by the homotetrameric association of a-subunits, encoded, in mammals, by the KCNMA1 gene^1^. This potassium channel is independently and allosterically activated by voltage and calcium^2,3^. This dual activation, together with its unique high conductance (ranging from 100 to 300 pS), make BK channel a key regulator of cell excitability, coupling intracellular Ca^2+^ signals to membrane potential^4^ over a wide dynamic range. The BK channel is widely expressed in many tissues, playing a role in smooth muscle tone maintenance, neurosecretion, and hearing acuity among others^1^.

Each of the BK channel alpha-subunits possesses a modular structure with 7 membrane spanning helices (S0-S6) and a large cytosolic domain with the tandem repeat of two Regulators of Conductance for K-like domains (RCK1 and RCK2). In the inner face of the membrane, the RCK domains associate in a ‘tetramer-of-dimers’ structure, commonly called the *gating ring*. Both RCK1 and RCK2 are where high affinity Ca^2+^ binding sites are located. The RCK1 domain contains the binding site with the lowest apparent affinity of the two (apparent affinities for the closed state, Kc, ≈ 26.8 μM and open state, K_o_, ≈ 5.6 μM)^5^. In this site, the divalent cation is coordinated in the human BK channel by the side chains of aspartate 367 and glutamate 535 (D367 and E535) and the backbone carbonyl oxygens of serines 533 and 600 and arginine 514 (S533, S600 and R514)^6,7^. In RCK2, the highest affinity coordination site - the *calcium bowl* - (K_c_≈3.1 μM/Ko≈0.88 μM)^5^ and the Ca^2+^ ions are coordinated by side chains of aspartates 895 and 897 (D895 and D897) and backbone carbonyl group of glutamine 889 and aspartate 892 (Q889 and D892) ^6–8^. Additionally, previous structural and functional studies have shown that the oxygen atom in the side chain of asparagine 449, (which comes from the RCK1 of the adjacent subunit), also coordinates Ca^2+^ in this binding site^6,7,9^. The mutation to alanine of N449 reduces the overall Ca^2+^ sensitivity^9,10^.

As a functional tetramer of RCK dimers, BK channel would be expected to have several interfaces of interaction, intra- and intersubunit. These assembly interfaces are crucial for the coordinated and synergistic BK channel activation. CryoEM full-length structures of the channel ^6,7,11^ unveiled some of these assembly interfaces of the gating ring and the potential key residues involved. In these works, two intersubunit assembly interfaces in the gating ring were shown as relevant for the activation mechanisms of the channel. The adjacent subunits interact with each other in two clear points: one RCK1-RCK2’ interface, with the N449 residue from RCK1 pointing towards the *calcium bowl* of the adjacent RCK2; and one RCK2-RCK2’ in the bottom part with electrostatic interaction of glutamate 955 and arginines 786 and 790 (E955, R786 and R790)^6,10^. Mutations in both interfaces were shown to dramatically affect Ca^2+^ sensitivity of the BK channel^10^. These two intersubunit interfaces mediate an interaction between adjacent monomers, evoking the concerted activation of the BK channel. Therefore, there is a dual role of these interfaces: Ca^2+^-binding facilitation and signal transduction to the pore.

To understand the dynamic movements of ion channels responsible for their activation states, and how the different domains of the proteins relate with each other, a variety of mutagenesis approaches have been employed. The development of unnatural amino acids (UAAs; also referred as non-canonical amino acids or ncAAs) provided a new dimension to this structure-function analysis studies by introducing new chemical groups beyond the natural reactive side chains. Photoactivatable amino acids have been widely employed in ion channel studies. These UAAs can form covalent bonds with nearby groups once they are irradiated with light (known as photocrosslinking). p-benzo-L-phenylalanine (BzF), together with L-azido-phenylalanine (AzF), are among the most common photoactivatable UAAs ^12–15^. Photoactivatable UAAs have assisted in the detailed study of protein function, identification of dimerization domains, study of desensitization and inactivation states of channels, mechanisms of allosteric modulators^16–18^. To date, BzF has been successfully incorporated in different ion channels to answer diverse biological questions^19^. In some of these studies, this tool has been shown useful to trap different conformations of the channel and elucidate their mechanisms of activation. For example, incorporation of BzF in the ligand-binding domain (LBD) of the AMPA receptor GluA2 inactivates the receptors following UV illumination^15^. On the other hand, the membrane domain of GluA2 was probed with AzF to delineate functionally-distinct regions^20^. The study of protein-protein interaction with BzF revealed state-dependent interactions and stoichiometry between KCNQ1 potassium channel and its auxiliary subunit KCNE1^21,22^. BzF incorporation in a linker present in ASIC channels disclosed the role of this domain in desensitization and recovery^23^. In the same channel, Braun et al. set a platform for high-throughput incorporation of BzF in hASIC1a and characterized the effects of different peptide activators^24^.

In this work we have introduced the BzF amino acid into the BK channel, and studied the dynamic interfaces between the different modules of activation of (voltage sensor, pore and gating domain and *gating-ring*. We incorporated BzF in 12 positions of the channel, recovering full-length and functional protein. Furthermore, we have identified Ca2+-dependent and voltage-independent current modification driven by UV light exposure at two different intersubunit RCK interfaces of the channel. Together, these results reinforce the idea that RCK intersubunit interfaces rearrange during Ca2+-mediated activation of the BK channel.

## Results and discussion

In order to functionally test the modules and residues of the BK channel involved in Ca^2+^ activation mechanisms, we aimed to insert the unnatural amino acid BzF in crucial domains of the protein. We generated a library of 14 BK channel variants where the *Amber* TAG codon was included at selected locations of the channel protein covering regions with particular functional relevance(figure 1). All mutations were generated on the background of BK667YFP and BK860YFP constructs, described previously^25–28^. Fusion of the fluorescent protein does not affect BK channel function and allowed us to quickly assess *Amber* suppression by BzF incorporation. BzF insertion sites covered five functional regions of the BK channel, the voltage sensor, the pore, the interface between the channel and the intracellular domains, and two interface regions in the intracellular domains that mediate Ca^2+^ activation (figure 1A).

**Figure 1.**
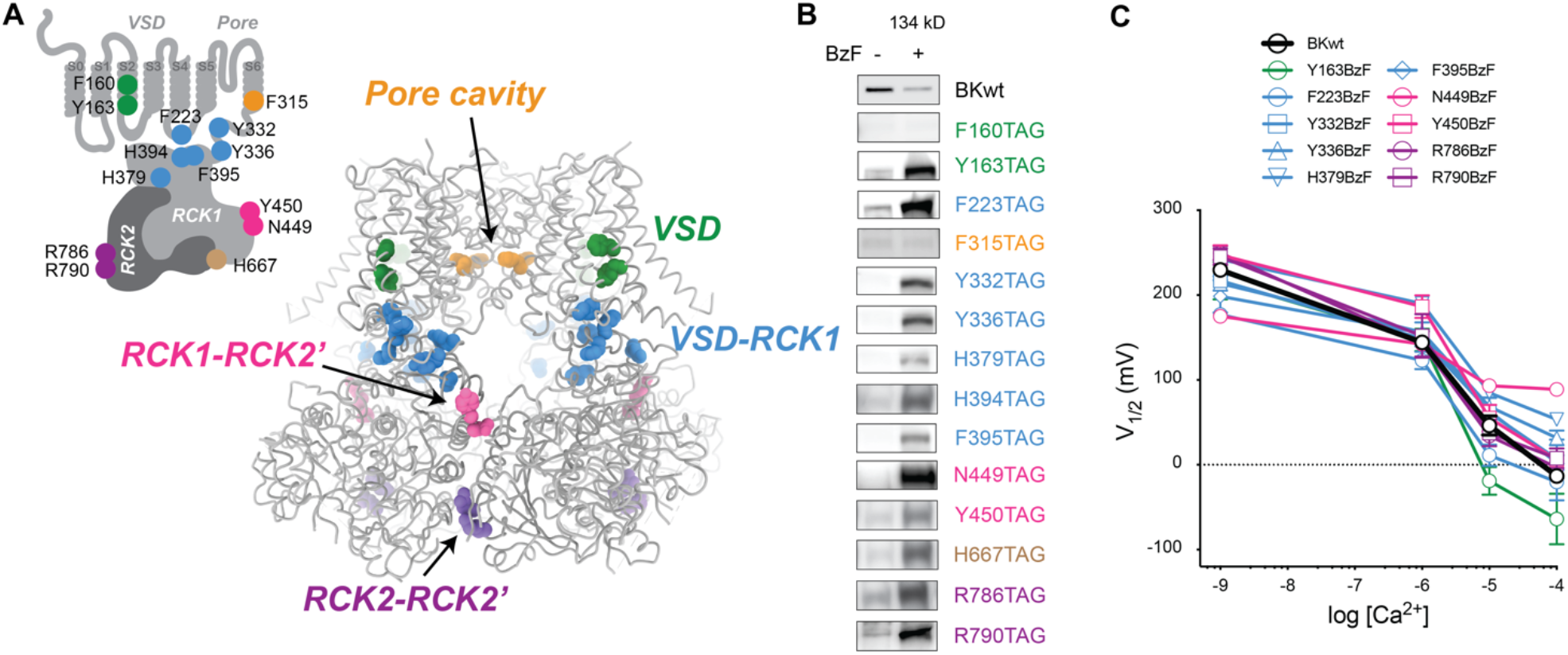
BK-BzF mutants rendered full-length proteins only under BzF incubated conditions. (A) Schematic diagram and structure location of the different positions selected for this study. The residues were selected along the channel to cover the different modules of the BK channel and their connections to the pore which could participate in the channel activation mechanisms: F160TAG and Y163TAG into the voltage-sensing domain (VSD); F223TAG, Y332TAG, Y336TAG, H379TAG, H394TAG and F395TAG to study Ca^2+^-binding sites to pore transduction pathways through the physical linker S6-RCK1 or through the VSD-RCK1 interface. F315TAG to alter the pore function without disturbing directly its permeability. N449TAG-Y450TAG and R786TAG-R790TAG were chosen to generate an intersubunit crosslinking between RCK domains in the different RCK1-RCK2’ and RCK2-RCK2’ interfaces respectively. H667TAG was used as a control because is a residue located in an unstructured region^7^ that would point outside the protein and in principle would not generate any effect under photocrosslinking (as fluorescent protein insertion did not affected channel function^25–28^) (B) Western-blot of the Amber suppression and full-length protein recovery in presence of BzF. BK mutants F160BzF and F315BzF did not show protein expression and were not further included in this study. (C) Mean log[Ca^2+^] vs V_1/2_ from the conductance-voltage curves fits with the Boltzmann equation for all of the mutants tested under 0, 1, 10 and 100 μM Ca^2+^.

### BK-TAG mutants incorporating BzF retained WT features

All but two BK-*TAG* mutants showed full-length protein recovery after incubation with BzF, showing specific and high yield incorporation of the photocrosslinker (figure 1B). All BK-BzF mutants produced voltage- and Ca^2+^-dependent outward K^+^ currents only in the presence of BzF in the medium (figure S1A). Figure 1C summarizes V_1/2_ vs log[Ca^2+^] relationships for WT BK and BK-BzF mutants. Interestingly, in some cases like the Y163BzF or F223BzF, the replacement of the original residues by a bulkier amino acid like BzF produced a dramatic effect in the slope of the G-V curve, as well as in their Ca^2+^ sensitivity (figure 1C, figure S1B and table 1). This strongly points towards a key residue and domain of the channel involved in the activation mechanisms.

**Table 1.**
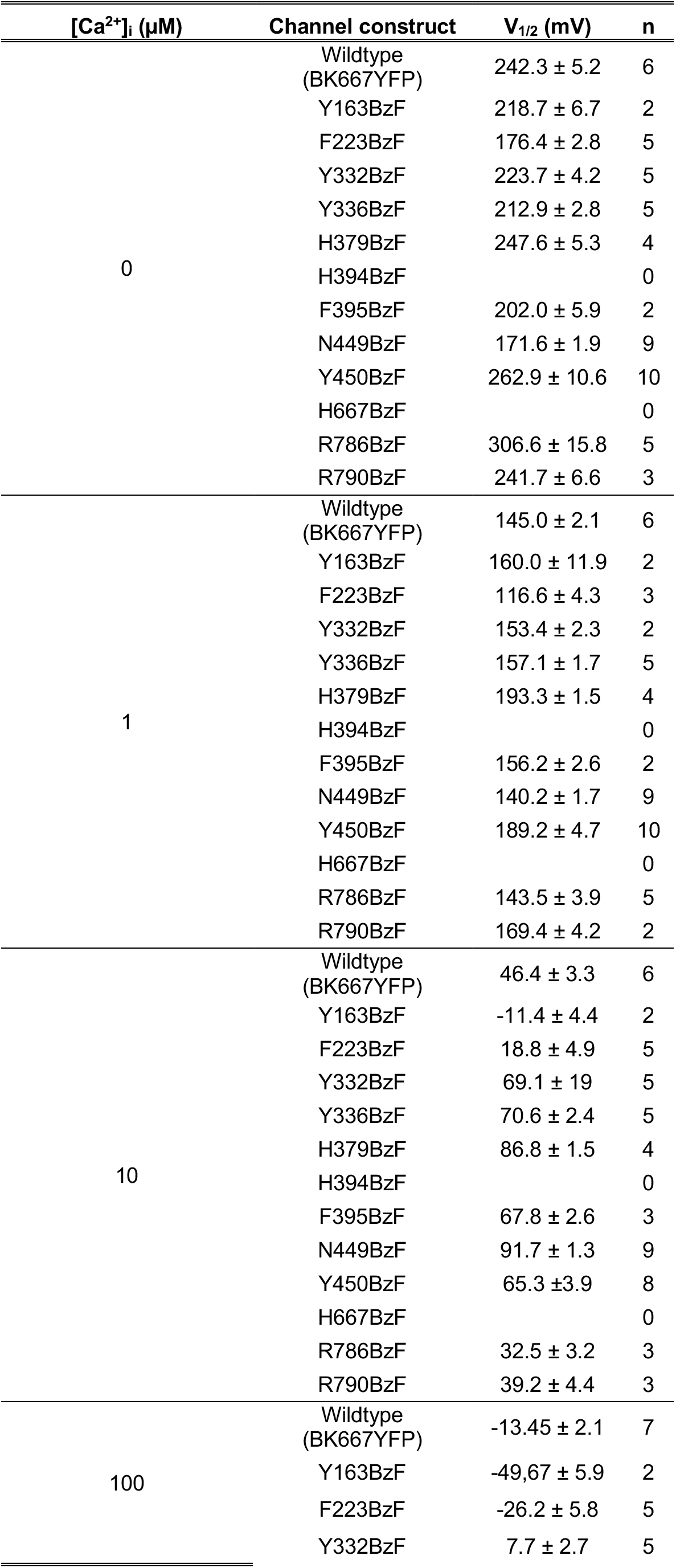
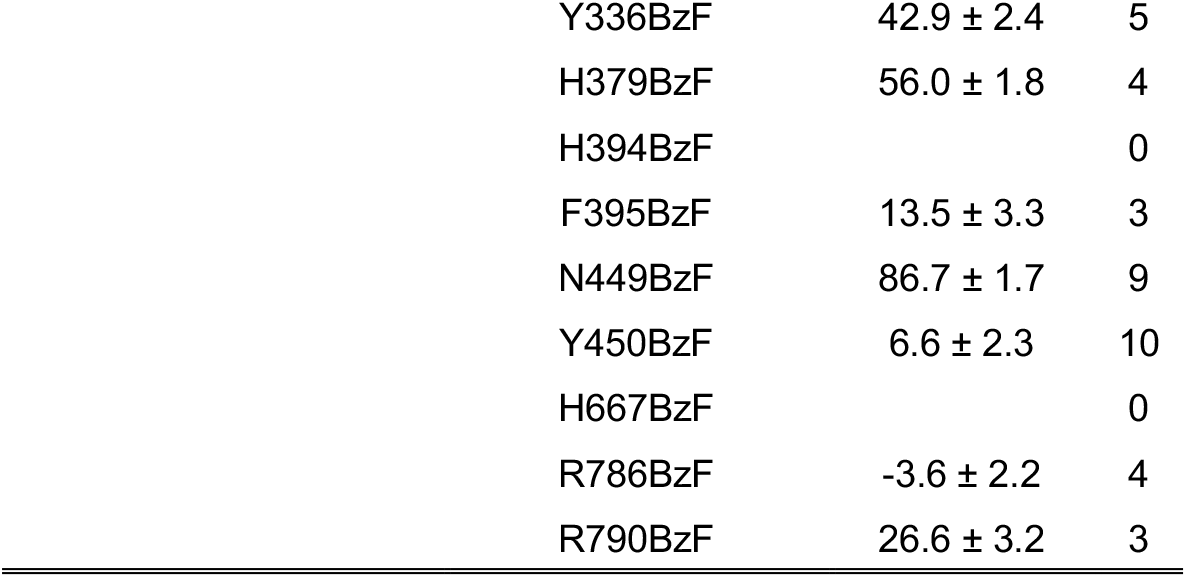
Boltzmann fit V_1/2_ for the WT and mutants under different Ca^2+^ concentrations. V_1/2_ is represented as mean ± SEM.

| $[\text{Ca}^{2+}]_i$ ($\mu\text{M}$) | Channel construct | $V_{1/2}$ (mV) | n |
| --- | --- | --- | --- |
| 0 | Wildtype (BK667YFP) | $242.3 \pm 5.2$ | 6 |
| | Y163BzF | $218.7 \pm 6.7$ | 2 |
| | F223BzF | $176.4 \pm 2.8$ | 5 |
| | Y332BzF | $223.7 \pm 4.2$ | 5 |
| | Y336BzF | $212.9 \pm 2.8$ | 5 |
| | H379BzF | $247.6 \pm 5.3$ | 4 |
|  | H394BzF |  | 0 |
| | F395BzF | $202.0 \pm 5.9$ | 2 |
| | N449BzF | $171.6 \pm 1.9$ | 9 |
| | Y450BzF | $262.9 \pm 10.6$ | 10 |
|  | H667BzF |  | 0 |
| | R786BzF | $306.6 \pm 15.8$ | 5 |
| | R790BzF | $241.7 \pm 6.6$ | 3 |
| 1 | Wildtype (BK667YFP) | $145.0 \pm 2.1$ | 6 |
| | Y163BzF | $160.0 \pm 11.9$ | 2 |
| | F223BzF | $116.6 \pm 4.3$ | 3 |
| | Y332BzF | $153.4 \pm 2.3$ | 2 |
| | Y336BzF | $157.1 \pm 1.7$ | 5 |
| | H379BzF | $193.3 \pm 1.5$ | 4 |
|  | H394BzF |  | 0 |
| | F395BzF | $156.2 \pm 2.6$ | 2 |
| | N449BzF | $140.2 \pm 1.7$ | 9 |
| | Y450BzF | $189.2 \pm 4.7$ | 10 |
|  | H667BzF |  | 0 |
| | R786BzF | $143.5 \pm 3.9$ | 5 |
| | R790BzF | $169.4 \pm 4.2$ | 2 |
| 10 | Wildtype (BK667YFP) | $46.4 \pm 3.3$ | 6 |
| | Y163BzF | $-11.4 \pm 4.4$ | 2 |
| | F223BzF | $18.8 \pm 4.9$ | 5 |
| | Y332BzF | $69.1 \pm 19$ | 5 |
| | Y336BzF | $70.6 \pm 2.4$ | 5 |
| | H379BzF | $86.8 \pm 1.5$ | 4 |
|  | H394BzF |  | 0 |
| | F395BzF | $67.8 \pm 2.6$ | 3 |
| | N449BzF | $91.7 \pm 1.3$ | 9 |
| | Y450BzF | $65.3 \pm 3.9$ | 8 |
|  | H667BzF |  | 0 |
| | R786BzF | $32.5 \pm 3.2$ | 3 |
| | R790BzF | $39.2 \pm 4.4$ | 3 |
| 100 | Wildtype (BK667YFP) | $-13.45 \pm 2.1$ | 7 |
| | Y163BzF | $-49.67 \pm 5.9$ | 2 |
| | F223BzF | $-26.2 \pm 5.8$ | 5 |
| | Y332BzF | $7.7 \pm 2.7$ | 5 |

|  |  |  |
| --- | --- | --- |
| Y336BzF | 42.9 ± 2.4 | 5 |
| H379BzF | 56.0 ± 1.8 | 4 |
| H394BzF |  | 0 |
| F395BzF | 13.5 ± 3.3 | 3 |
| N449BzF | 86.7 ± 1.7 | 9 |
| Y450BzF | 6.6 ± 2.3 | 10 |
| H667BzF |  | 0 |
| R786BzF | -3.6 ± 2.2 | 4 |
| R790BzF | 26.6 ± 3.2 | 3 |

### Photocrosslinking of cytosolic interfaces altered BK function

We hypothesized that photocrosslinking of BzF inserted at sites relevant for BK channel gating would result in UV-dependent alterations of channel function. Exposure to UV light altered gating behavior of 4 out of the 12 mutants generated (figure 2). Photocrosslinking of BzF inserted at residues N449 or Y450 induced a current reduction of 66% and 47% respectively, only at high Ca^2+^ concentrations (figure 2B). Contrarily, photocrosslinked R786 or R790 positions induced a potentiation of the channel current of 243% and 72% in the absence of Ca^2+^ (figure 2B). Interestingly, these residues form intersubunit interfaces in the BK channel gating ring^6,10^.

**Figure 2.**
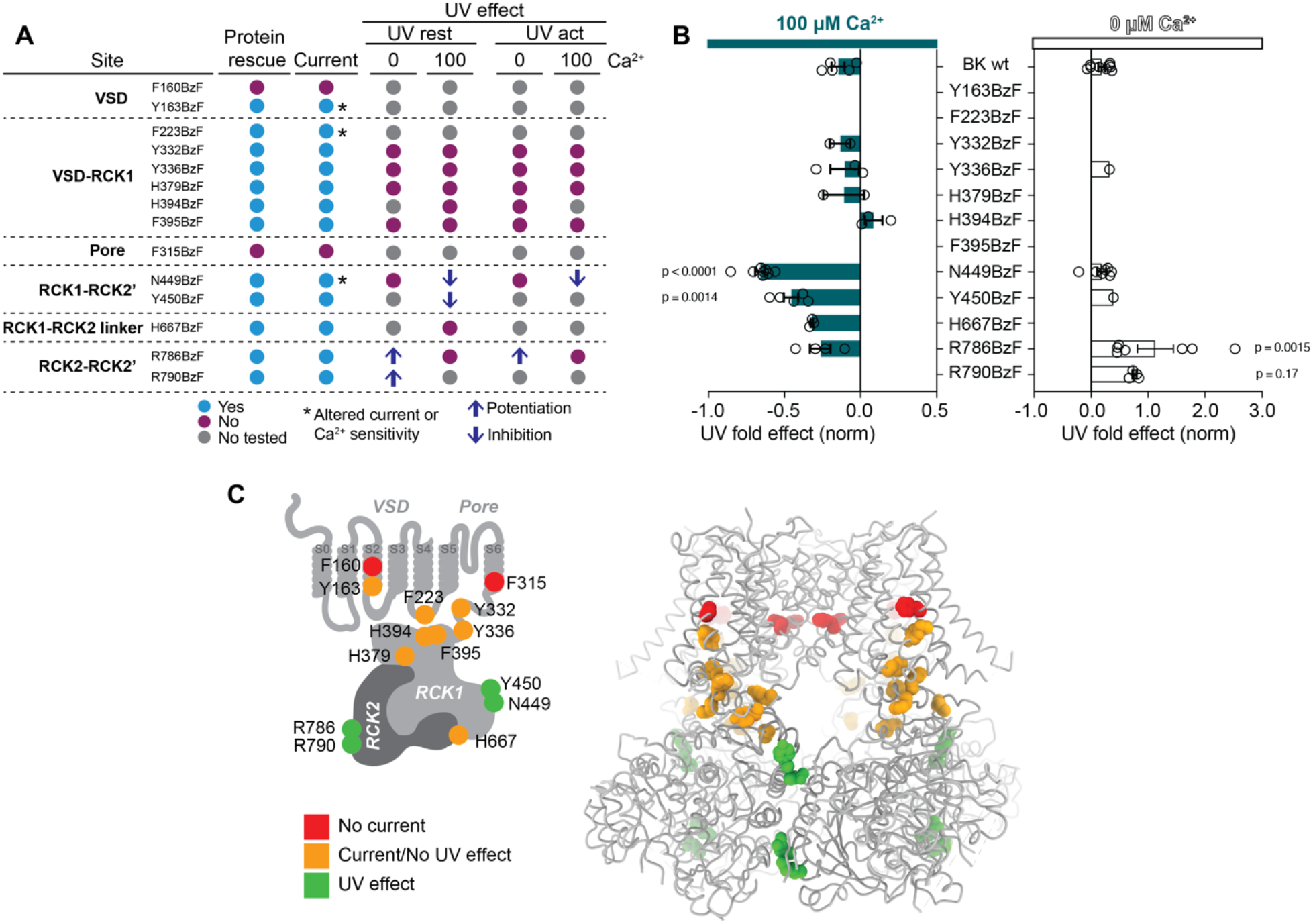
Summary of BK-BzF mutants function and UV-specific effects. (A) Summary table of mutants characterization. (B) UV fold effect in 0 and 100 μM Ca^2+^ for the conditions and mutants tested in this work (ANOVA, p-values from Dunett’s multiple comparison test). (C) Same schematic diagram and structure location of the residues as in figure 1 but color coded regarding the functional recovery and UV photocrosslinking effect.

Can the effects of photocrosslinking on channel gating be explained by alteration of the intersubunit interfaces? We compared the available human full-length structures for Ca^2+^ free and Ca^2+^ bound BK channels (figure 3; PDB: 6V3G and 6V38^7^). The BK channel presents a domain-swapped structure so that the VSD of each subunit is positioned above f the RCK domains of the adjacent subunit. The individual RCK domains interact with each other at different points. It has been proposed that the full-length BK channel an RCK1 and RCK2 Ca^2+^ binding sites are connected by an intrasubunit bridge^6,11^, which is relevant for channel activation by Ca^2+ 29^. In addition, the structure showed RCK1 and RCK2 intersubunit interactions at two regions: one is the RCK1-RCK2’ interaction involving the N449 residue that coordinates the Ca^2+^ of the adjacent subunit RCK2 *calcium bowl* (here designated as site A, figure 3A and 3B); the other interaction occurs between RCK2-RCK2’ involving the electrostatic interaction between residues R786, R790 and E955 (site B; figure 3A and 3B).

**Figure 3.**
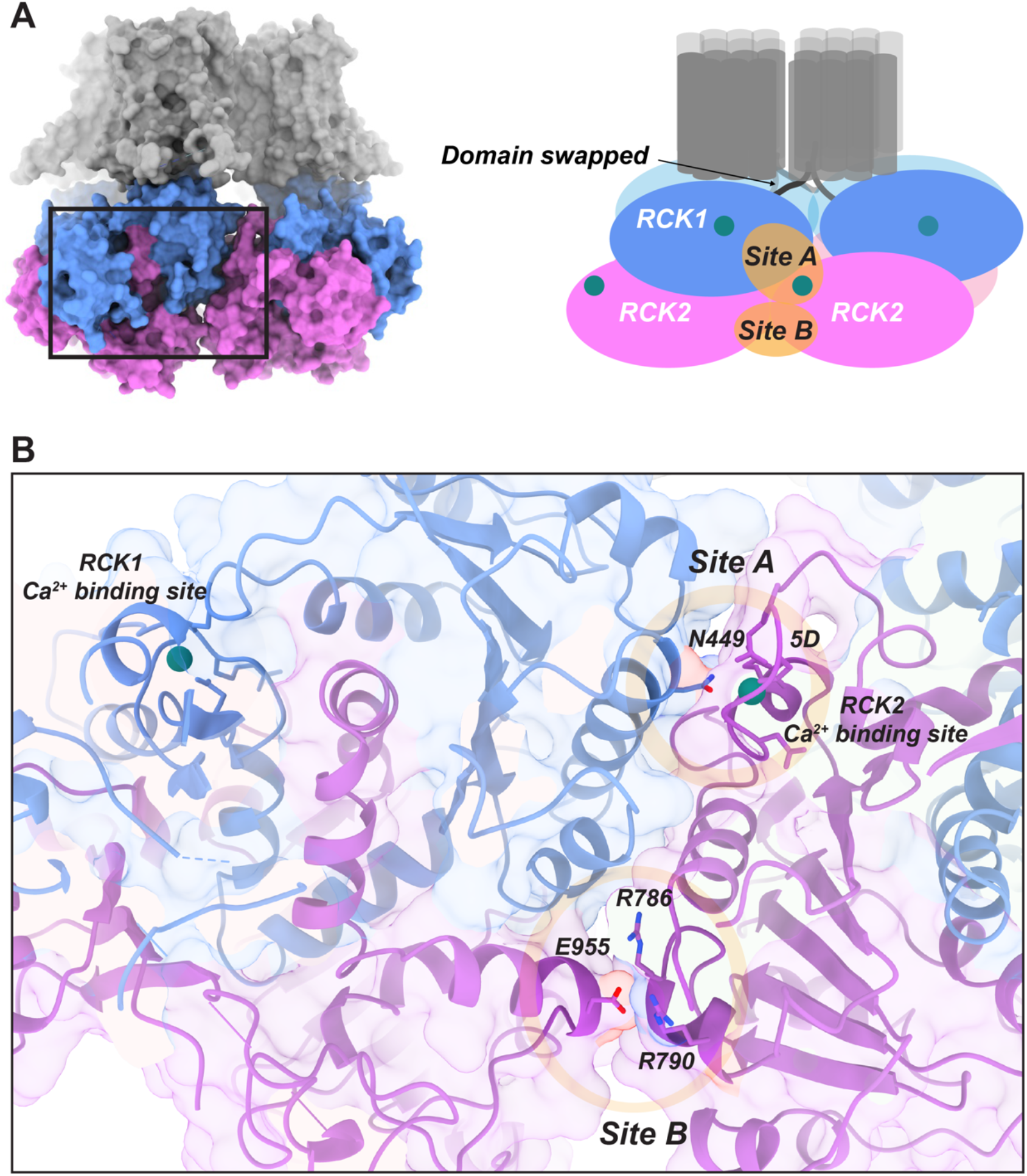
Intersubunit interfaces of the *gating ring* of the BK channel. (A) Surface depiction of homotetramer assembly of human BK channel structure (PDB: 6V38; grey: transmembrane domains, blue: RCK1, pink: RCK2) and diagram of the structure of the channel. Homotetramer configuration presents a domain swap structure where the cytosolic domain of one monomer is below the transmembrane domain of the adjacent subunit. This cytosolic domain presents two clear intersubunit interfaces: RCK1-RCK2 (Site A) and RCK2-RCK2 (Site B). (B) Residues involved in the intersubunit interactions and Ca^2+^ binding sites. Site A is constituted by N449 and its coordination of the Ca^2+^ ion of the RCK2 Ca^2+^ binding site. This site is far from its own RCK1 Ca^2+^-binding site. Site B is further from Ca^2+^ binding sites and constitutes a RCK2-RCK2 interaction on the bottom part of the channel.

These two interaction interfaces do not move significantly when Ca^2+^ binds to the channel (figure 3B). However, they seem to be pivotal residues around which the RCK domains rotate in the absence or presence of Ca^2+^. It is important to remark that these structures were obtained in the absence of membrane potential, most likely precluding the structure to fully reflect the structural features of the allosterically coupled Ca^2+^ and voltage activation of the channel.

### BzF insertion at RCK intersubunit interfaces produced a Ca^2+^-dependent current reduction or potentiation under UV exposure

We interrogated the role of intersubunit interfaces (site A and B; figure 3) in BK channel activation by Ca^2+^ with the use of photocrosslinking. A first approach consisted of studying the effect of applying UV light in the absence or presence of saturating 100 μM Ca^2+^ while applying hyperpolarizing or depolarizing voltages (figure 4A-D). In the absence of Ca^2+^, photocrosslinking of the N449BzF mutant (site A) did not show any significant effect when UV light was applied during the hyperpolarizing or the depolarizing pulse (figure 4E and G). However, under saturating Ca^2+^ conditions, and independently of voltage, a current reduction of 55-60% was observed in this mutant (figure 4F and H). This effect can be attributed to specific photocrosslinking of the 449 site, since BK667YFP channels did not show significant UV-dependent alteration of current levels (figure S4-1B and D). Consistent with this idea, BzF insertion into an adjacent residue, Y450BzF, showed similar current reduction at saturating Ca^2+^ conditions (figure S4-1E), which was not observed in other regions of the channel (e.g. H394BzF, figure S4-1F). Current decay from individual N449BzF and Y450BzF experiments can be fitted to a single exponential decay (table 2).

**Figure 4.**
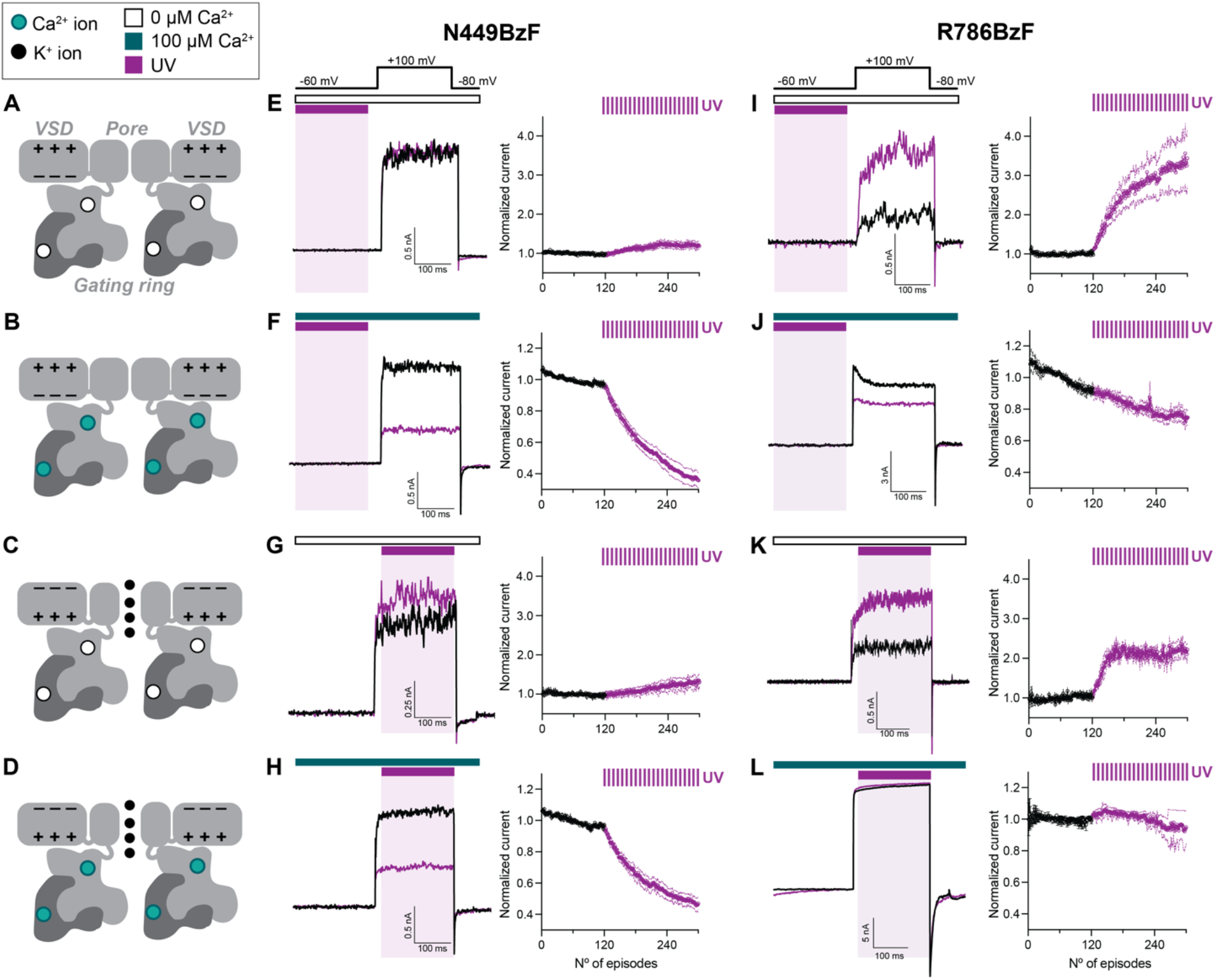
Photocrosslinking in site A and site B modify BK channel current in opposite directions in a voltage independent and a Ca^2+^ dependent manner. (A-D) Diagrams of a BK channel in the different states where the experiments were performed. (E-H) Effects of N449BzF mutant with the UV applied under different conditions. Only in the presence of saturating Ca^2+^ (100 μM), N449BzF showed a UV-dependent current decay both in hyperpolarized and depolarized patches. Left panels show two representative traces at the beginning and after UV exposure of the same patch. Right panels show the averaged and normalized kymograms for the (I-L) In the case of the R786BzF, UV applied in the absence of Ca^2+^ produced a potentiation that was not observed when saturating 100 μM Ca^2+^ was present.

**Table 2.** Exponential fit parameters and UV exposure time for the photocrosslinking.

| Mutant | UV application potential (mV) | [Ca<sup>2+</sup>]<sub>i</sub> (μM) | n | Crosslinking effect | t (episodes) | Cumulative UV exposure at t (s) | Plateau |
| --- | --- | --- | --- | --- | --- | --- | --- |
| BK wt (667YFP) | -60 | 0 | 9 | - |  |  |  |
|  | -60 | 100 | 5 | - |  |  |  |
| N449BzF | -60 | 0 | 8 | - |  |  |  |
|  | 100 | 0 | 3 | - |  |  |  |
|  | -60 | 1 | 4 | Current reduction | 229 | 45.8 | 0.44 |
|  | -60 | 10 | 2 | Current reduction | 149 | 29.8 | 0.46 |
|  | -60 | 100 | 8 | Current reduction | 93 | 18.6 | 0.26 |
|  | 100 | 100 | 5 | Current reduction | 108 | 21.6 | 0.35 |
| N449BzF + D367A | -60 | 100 | 4 | - |  |  |  |
| N449BzF + 5D5A | -60 | 100 | 3 | Current reduction | 219 | 43.8 | 0.4 |
| N449BzF + D367A + 5D5A | -60 | 100 | 5 | - |  |  |  |
| Y450BzF | -60 | 100 | 5 | Current reduction | 196 | 39.2 | 0.39 |
| R786BzF | -60 | 0 | 7 | Current potentiation | 27 | 5.4 | 2.2 |
|  | 100 | 0 | 4 | Current potentiation | 17 | 3.4 | 2.2 |
|  | -60 | 1 | 2 | - |  |  |  |
|  | -60 | 10 | 2 | - |  |  |  |
|  | -60 | 100 | 6 | - |  |  |  |
|  | 100 | 100 | 3 | - |  |  |  |
| R786BzF + D367A | -60 | 100 | 0 |  |  |  |  |
| R786BzF + 5D5A | -60 | 100 | 0 |  |  |  |  |
| R786BzF + D367A + 5D5A | -60 | 100 | 0 |  |  |  |  |
| R790BzF | -60 | 0 | 4 | Current potentiation | 146 | 29.2 | 2.3 |

We next explored the effects of photocrosslinking at site B interface. In contrast with site A results, R786BzF showed a potentiation of the current in the absence of Ca^2+^ (figure 4I and K). Current levels reached up to 3.5 fold when UV was applied during the hyperpolarizing pulse (figure 4I) and 2 fold when UV was applied while the voltage sensors were activated by depolarization (figure 4K). In the presence of saturating Ca2+, no effect of UV was observed, consistent with larger distances between photocrosslinking partners than in the absence of Ca^2+^ (figure 4J and L). Interestingly, UV independent rundown of the current was consistently observed for this mutant (figure 4J). We observed again that the UV-dependent potentiation is specific to the site B interface and absent in the wildtype channel (figure S4-1A and D). Photocrosslinking of adjacent residue R790BzF also showed potentiation of about 2 fold while other residues along the channel did not have any effect (i.e. R790BzF and Y336BzF; figure S4-1G-H).

The specificity of our results was further confirmed by the observation that the intensity of the UV-dependent effects is not further increased when UV light application is stopped (figure S4-2I-J). In addition,the effect was only detected under the specific Ca^2+^ conditions tested, while immediate application of different Ca^2+^ concentrations (0 μM for N449BzF and 100 μM for R786BzF) blunted the photocrosslinking effect (figure S4K-L). Altogether, these results corroborate that the observed effects are state-dependent, only occurring at specific conformations at which the distance between the inserted BzF residue and its photocrosslinking partner is close enough for recombination.

Together, these results provide information about key activation rearrangements of the channel. In the absence of Ca2+, RCK2 domains from adjacent subunits are closer at the site A region. The fact that locking this conformation by photocrosslinking potentiates activation of the channel could be attributed to the facilitation of channel opening by voltage. This could be explained by two possible mechanisms: 1) a negative coupling between VSD and RCK2, or 2) a locked conformation in an intermediate step of activation that expands the gate of the channel, changing its unitary conductance. On the other hand, the current reduction observed when photocrosslinking site B interaction can be explained by a decoupling of the VSD to the gating ring. Alternatively, this effect could be due to the inability of the interface to further rotate and activate the channel.

### RCK1 Ca^2+^ binding site occupancy underlies the N449BzF photocrosslinking effect

We next interrogated if the observed Ca^2+^-dependent photocrosslinking effects on channel function could be attributed to activation of specific Ca^2+^ binding sites. We first tested ourphotocrosslinking protocols under four different Ca^2+^ concentrations (figure 5A-C). In the case of N449BzF, the effect clearly showed Ca^2+^ dose-dependent effect oncurrent reduction (figure 5A). While the kinetics for the current reduction were faster for 10 μM than for 1 μM Ca^2+^ (tau= 149 vs 229 episodes), the difference of the total effect between these two intermediate calcium concentrations was not significant (plateau 0.46 vs 0.44 respectively). Because the affinities of both Ca^2+^ binding sites differ at least in one order of magnitude and the high affinity RCK2 binding site is saturated below 10 μM Ca^2+^, these results led us to propose that the lower affinity RCK1 binding site needs to be occupied by Ca^2+^ for the channel to complete photocrosslinking.

**Figure 5.**
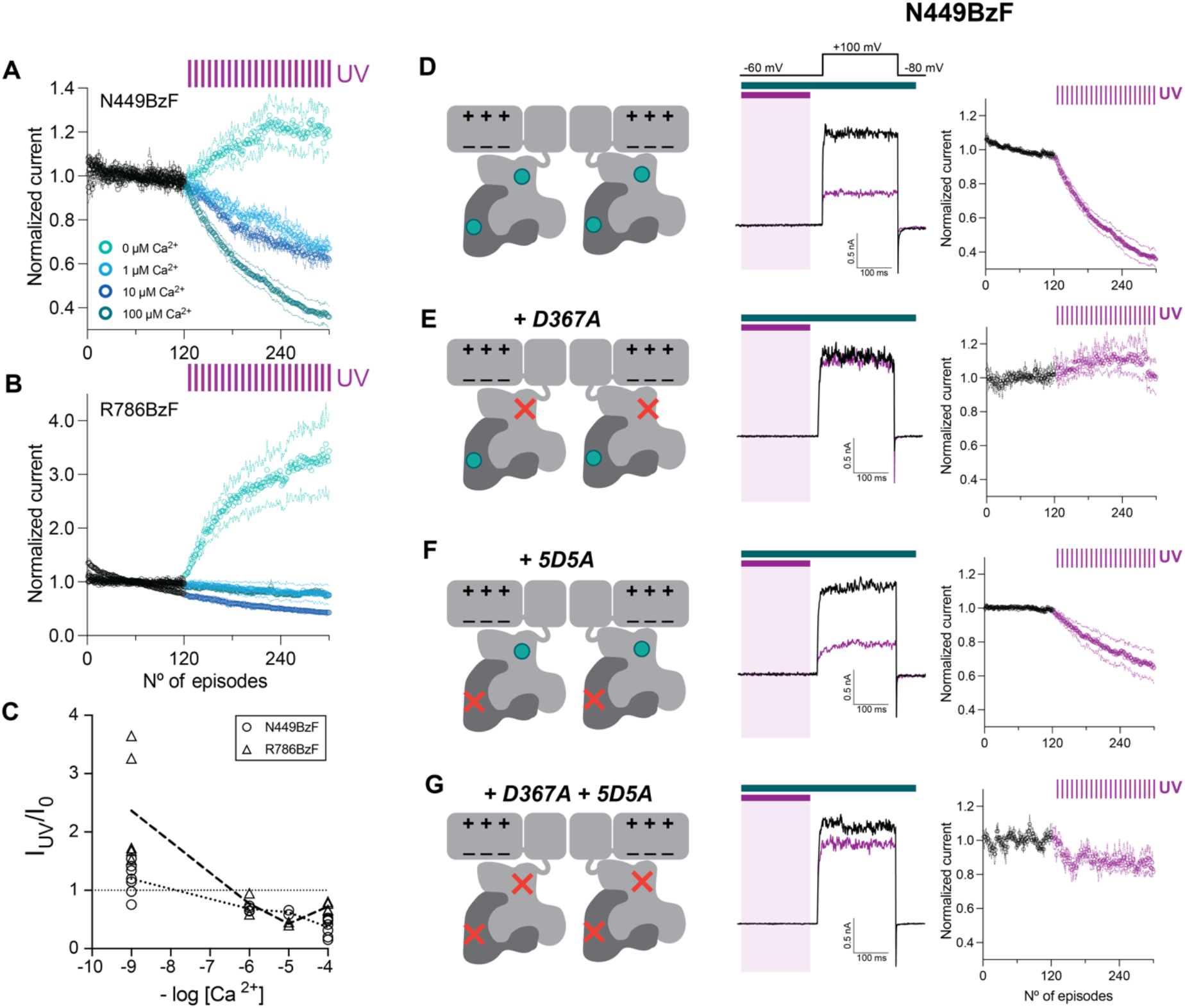
Prevention of Ca^2+^ binding to the RCK1 site abolished the N449BzF effect. (A) N449BzF mutant showed a Ca^2+^ concentration dependent current decay. The ‘potentiation’ of 0 μM Ca^2+^ is not specific and was similarly observed in the WT (figure S4-2) and it is significantly different for the one of the R786BzF mutant. (B) For the R786BzF, only in the absence of Ca2+, the potentiation effect was observed. The minimum application of Ca2+, 1 μM, did not show any potentiation (nor 10 or 100 μM either). (C) Summary of the individual experiments at the different Ca^2+^ against the log[Ca2+]. (D-G) Effect of the different mutants under UV light. D367A, a mutant that abolishes Ca^2+^ binding to the RCK1 site, abolished the current reduction effect while 5D5A, the equivalent preventing Ca^2+^ binding mutant of the RCK2 site, still showed a UV-dependent current reduction. The occupancy of RCK1 site by Ca^2+^ ions seems to be crucial for the localization of N449BzF close enough to the photocrosslinking partner.

We tested this hypothesisby using mutations that are well known to abolish Ca^2+^ binding to specific binding sites. Mutation of residue 367 to Alanine (D367A) has been described to abolish the function of the RCK1 binding site. In the background of the N449BzF construct, the D367A mutation completely abolished the effect of photocrosslinking on BK currents (figure 5E). In contrasts, mutation of the RCK2 high affinity calcium bowl (5D5A) a proportion of the UV-dependent current reduction remained (figure 5F). Full abrogation of Ca^2+^ binding to the channel by introducing thedouble mutation (figure 5G) rendered effects that were comparable to mutation of the RCK1 sites, leading us to conclude that occupancy of the RCK1 Ca^2+^ binding site is essential for the photosensible current reduction to occur. Together, these findings point towards a mechanism where N449 and the neighboring residues involved in the RCK1-RCK2 intersubunit interface position close to the RCK2 binding site only when RCK1 binding site is occupied by Ca^2+^. The fact that N449, being far from the RCK1 binding site, needs its occupancy to drive the effect and be closer to the RCK2 binding site of the adjacent subunit suggests a form of cooperativity between both Ca^2+^ binding sites that it will take place once both binding sites are occupied.

### UV-dependent potentiation of R786BzF channel function is only observed in the absence of Ca^2+^

The effect of the site B intersubunit interface was studied using the R786BzF constructs. In contrast with site A interactions, UV exposure of R786BzF leads to current potentiation in the absence of Ca^2+^. Application of a low concentration of Ca^2+^ 1 μM which, in principle, would only saturate the RCK2 site, did not show the potentiation effect (figure 5B-C). Because the effect was seen only in the absence of Ca^2+^, we did not pursue a binding sites mutagenesis analysis like in the N449BzF. This result led us to conclude that the RCK2-RCK2 intersubunit close interaction is only seen in the absence of Ca^2+^. Once Ca^2+^ is bound to the high affinity Ca^2+^ binding site, the distance between R786BzF and the adjacent RCK2 is not enough for an effective crosslinking, indicating a separation of the interfaces.

### In silico analysis of the distances between residues BzF reactive group and neighbor alpha carbons identified plausible intersubunit crosslinking

One of the limitations of the use of photocrosslinkers is the lack of control of the crosslinking partner within the channel structure. To explore whether any residues could be close enough to undergo photocrosslinking, we performed an in silico analysis using the published human structures of the BK channel^7^. We mutated the N449 to BzF using the SwissSideChain database and selecting the rotamer with the highest probability. Afterwards, we measured the distances between the carbon from the ketone reactive group of BzF and the Ca carbons closer to it (from RCK2’ calcium bowl region) (figure 6A1). A comparison between the Ca^2+^ free and Ca^2+^ bound structures showed a clear hotspot for photocrosslinking around Q889 residue (figure 6A2), being the only point where a distance slightly below the maximum ideal 3.1Å is reached^30^.

**Figure 6.**
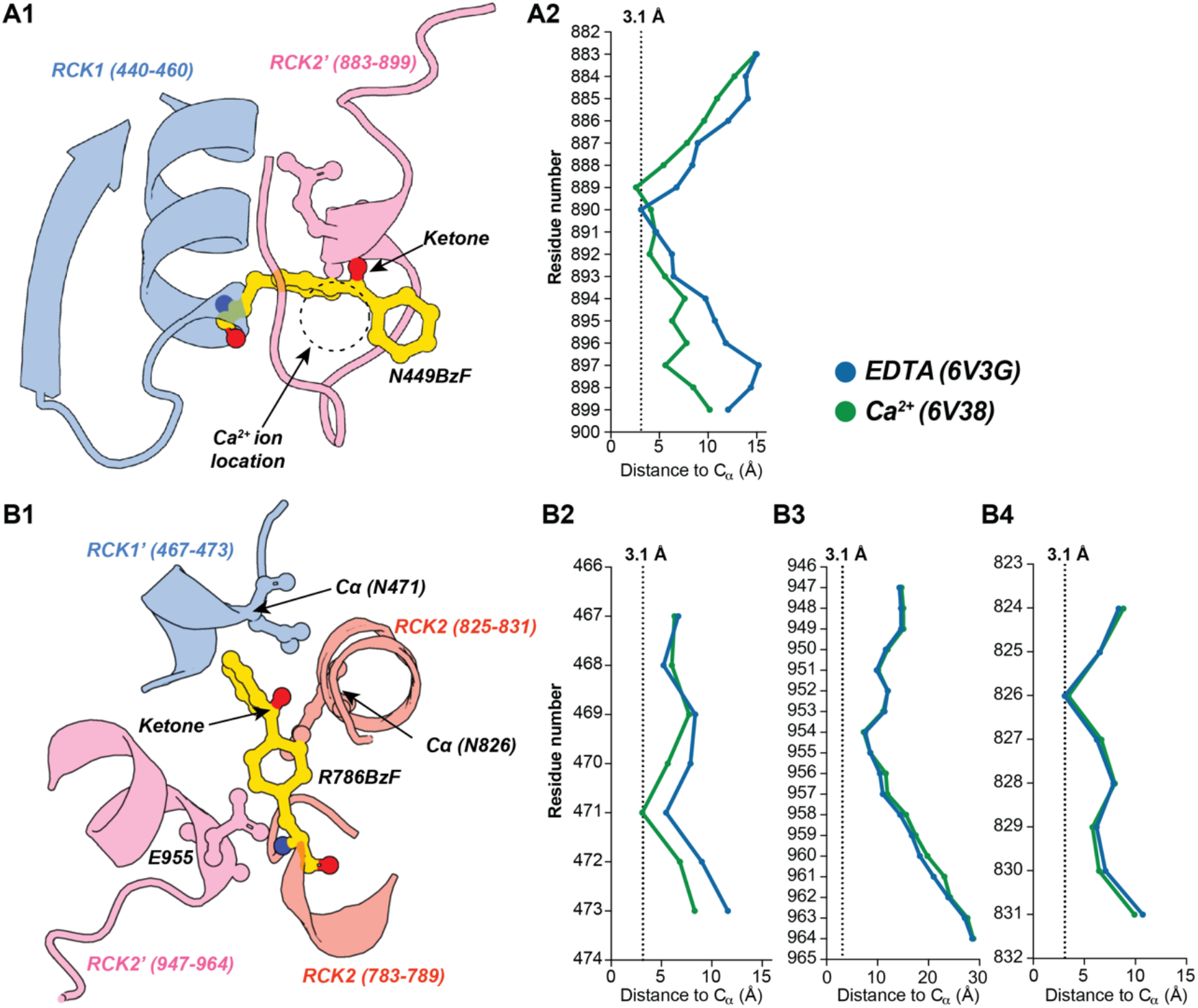
*In silico* analysis of the potential distances for photocrosslinking showed a hotspot in the N449BzF but not in R786BzF. Distance calculation between the ketone active group of the benzophenone mutagenized in silico (using SwissSideChain database) in the Ca^2+^ bound or Ca^2+^ free structures (PDB: 6V38 and 6V3G respectively) and the alpha carbon of the closest environment. (A1) For the N449BzF, the benzophenone insertion invades the Ca^2+^ binding site of the RCK2 subunit (which explains an inability of the Ca^2+^ to bind the site once the photocrosslinking happens. (A2) the Ca^2+^ bound structure showed a hotspot of photocrosslinking that changes between both structures and matches with the observed effect in 100 μM Ca2+. We took 3.1Å as the maximum distance for photocrosslinking^30^. (B1) In the case of R786BzF, the environment is more complicated, with certain residues close from different domains of the protein. In this case we could not determine a plausible hotspot for photocrosslinking that explains our potentiation effect.

BzF inserted in R786BzF has three different domains close enough to photocrosslink. Surprisingly, the furthest one was the loop where E955 is located (figure 6B1,B3) and which forms part of the RCK2-RCK2’ interface. The other close residues to photocrosslink are the N471 on the RCK1’ (figure 6B1,B2) and the N826 in the same RCK2 (figure 6B1,B4). Interestingly, the only one that suits the situation of 0 μM Ca^2+^ and below 3.1 Å is the N826. We have to keep in mind that this analysis is based on the most probable rotamer by using the ‘Rotamer’ tool of Chimera software which has some limitations regarding the local reorganizations in the backbone by insertion of the mutated amino acid, only considering the side chain. This analysis was useful to explore the environment surrounding the R786 position, however, in this case we could not extract more information.

## Materials and methods

### Molecular biology, Amber codon mutagenesis, and plasmid purification

The full-length human BK channel (GenBank accession number U11058) fluorescent derivatives BK667YFP and BK860YFP were selected from a library previously available in the lab^25^. All the inserts were in a mammalian expression with an ampicillin resistance vector under a CMV promoter (pBNJ13). Plasmids expressing the orthogonal pair of BzF specific aminoacyl-tRNA synthetase (aaRS) and Amber-supressor tRNA were kindly gifted by Dr. Thomas Sakmar (Rockefeller University, New York, USA). These constructs were previously designed and validated to be used in mammalian cell lines^31^. *Amber* codon mutagenesis in the selected positions were performed using the QuickChange Lightning Site-directed Mutagenesis kit (Agilent Technologies, California, USA) following the manufacturer protocol. XL-10 Gold Ultracompetent cells (Agilent Technologies, California, USA) were for plasmid transformation, growth, storage and purification. Plasmid purifications for mammalian cell lines transfection were performed using high-yield column-based commercial NucleoBond Xtra Midi kit (Macherey-Nagel, Germany) following usage protocols.

### Cell culture, transfection and UAA incorporation

HEK293T cells were grown in complete DMEM. DMEM High-glucose (4500 mg/l), with stable-glutamine and Sodium pyruvate (Biowest, Nuaillé, France) was supplemented with 10% fetal bovine serum (FBS) (Biowest, Nuaillé, France) and 1% penicillin/streptomycin (Biowest, Nuaillé, France). Cells were chemically transfected with jetPrime (PolyplusTM, Strasbourg, France) following manufacturer recommendations. The transfection ratios for plasmids expressing BK-TAG: tRNA^CUA^: aaRS were optimized and fixed to 1:1:1. Culture media with transfection reagent was replaced with fresh media between 4-6 hours after transfection. For the UAA incorporation, p-benzoyl-l-phenylalanine supplemented complete DMEM was freshly made before media change. For 15 mL of DMEM supplemented with 1 mM BzF, 4 mg of BzF (MW: 269.3; Bachem, Bubendorf, Switzerland) were dissolved in 300 μL of HCl (1M). The dissolved BzF was immediately added to 15 mL of 37 °C pre-warmed complete DMEM and pH was adjusted to 7.4 with NaOH. This standard protocol was extensively used and optimized before ^15,20,31,32^.

For electrophysiology experiments, cells were seeded in Ø13 mm coverslips precoated with poly-L-lysine (Sigma Aldrich). For biochemistry experiments, cells were grown in Ø100mm pre-treated culture dishes (Nunclon, Thermo Scientific).

### Protein biochemistry, SDS-PAGE and western blot

To check specific UAA insertion and full-length protein recovery, cells were collected in PBS after 48 hours post-transfection. After pelleting by centrifugation, cells were resuspended in TENT lysis buffer (in mM: 50 Tris-HCl, 150 NaCl, 5 EDTA and 1% Triton X-100) supplemented with cOmplete™ Mini-EDTA free protease inhibitor cocktail (Roche, Germany). Lysates were incubated 5 min on ice and centrifuged 10 min - 14000 x g at 4°C. Supernatant was kept on ice, quantified with bicinchoninic acid assay (BCA) and Laemmli loading buffer was added. SDS-PAGE was performed in Mini-PROTEAN® TGX Stain-Free™ Precast gels (10%) (BioRad, USA). Proteins were transferred to a PVDF membrane using the semidry system Trans-Blot® Turbo™ Transfer Starter System (PVDF) for Western Blot analysis (BioRad, California). Full-length BK channel was detected using Anti-Maxi Potassium channel alpha/SLO primary antibody [L6/60] (ab192759; Abcam, UK) and a goat anti-mouse HRP conjugated (P0447, Dako, Denmark).

### Electrophysiology recordings and BzF photoactivation

Inside-out patch clamp recordings were performed 48-72 hrs post-transfection with an Axopatch 200B amplifier and a Digidata 1550A digitizer (Molecular Devices, USA). Digitizer was controlled through a conventional PC. Fluorescent cells were illuminated with a multicolor Spectra X LED engine (Lumencor, Oregon, USA) and captured under a Nikon Eclipse Ti-U (Nikon, Japan) inverted microscope with an emCCD camera (iXon Ultra 888, Oxford Instruments Andor, UK). Microscope LED, and camera were synchronized and controlled using μManager (Image J, USA) with a Z440 workstation (Hewlett-Packard, California, USA).

BK channel currents were recorded under symmetrical potassium conditions with a pipette solution containing, in mM: 80 KMeSO3, 60 N-methylglucamine-MeSO3, 20 HEPES, 2 KCl, and 2 MgCl2 (pH 7.4); and a bath solution containing (in mM) 80 KMeSO3, 60 N-methylglucamine-MeSO3, 20 HEPES, 2 KCl. 1 mM of Ca^2+^ chelator hydroxyethyl ethylenediamine triacetic acid (HEDTA) was added to all the Ca^2+^ concentrations below 100 μM free Ca^2+^. The amount of CaCl2 added to obtain the desired free Ca^2+^ concentration was calculated with MaxChelator (Bers et al., 1994; downloaded from maxchelator.stanford.edu). Ca^2+^ free concentrations were corroborated using a Ca^2+^ electrode (Orion, Thermo Fisher Scientific, Massachusetts, USA)

Photoactivation of the UAAs was done with a 365 nm wavelength UV LED (ThorLabs, USA) remotely controlled through the electrophysiology computer and digitizer with TTL pulses. Dichroic mirror T400lp (Chroma Technology, Vermont, USA) mounted on a Nikon adapter filter cube was used for total UV-LED light reflection to the focal plane.

### Data analysis and software

Current data was firstly analyzed in Clampfit 11 (pCLAMP11, Molecular Devices, USA) to measure current traces (peaks and averages). Results were exported to Excel (Microsoft, USA) and Prism (GraphPad, California, USA), where plot representations and statistical analysis were performed.

Conductance versus voltage (G-V) analysis was obtained by normalizing tail current value at - 80 mV (I) to the highest current value (Imax) obtained under the maximum Ca^2+^ concentration condition (mainly 100 μM Ca^2+^) and represented versus voltage. Curves were generated from tail current amplitudes normalized. The resultant curves were fitted with the Boltzmann equation:

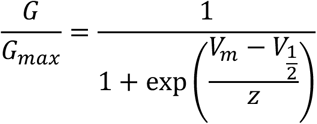

where, V_1/2_ is the voltage of the half-maximum activation, z is the slope of the curve, Vm is the test potential and Gmax is the maximal conductance.

To evaluate the effect of photocrosslinking in the overall function of the BK channel, different protocols of UV application and recording were applied, as indicated in the results. Due to the limited availability of the unnatural amino acid (fixed number of channels per patch), the expected effect of its photocrosslinking should describe an exponential effect of current reduction or potentiation. The photocrosslinking effects were measured as a change in the steady-state current at the end of an activating depolarizing pulse of +100 mV along the cumulative exposure to UV. For all the mutants which described this exponential effect (decrease or increase), a single exponential was fitted to the UV exposed episodes as previously reported (Ding and Horn, 2001). The current was normalized to the average of the baseline (before UV application) for comparison among patches (*I_t_ /I_baseline_*).

For protein visualization, in silico unnatural amino acid mutagenesis (with SwissSideChain plugin; Gfeller et al., 2013) and atom-to-atom distances calculation, we used Chimera software, from Resource for Biocomputing, Visualization, and Informatics at the University of California, San Francisco (supported by NIH P41 RR-01081).

## Supplemental figures

**Figure S1.**
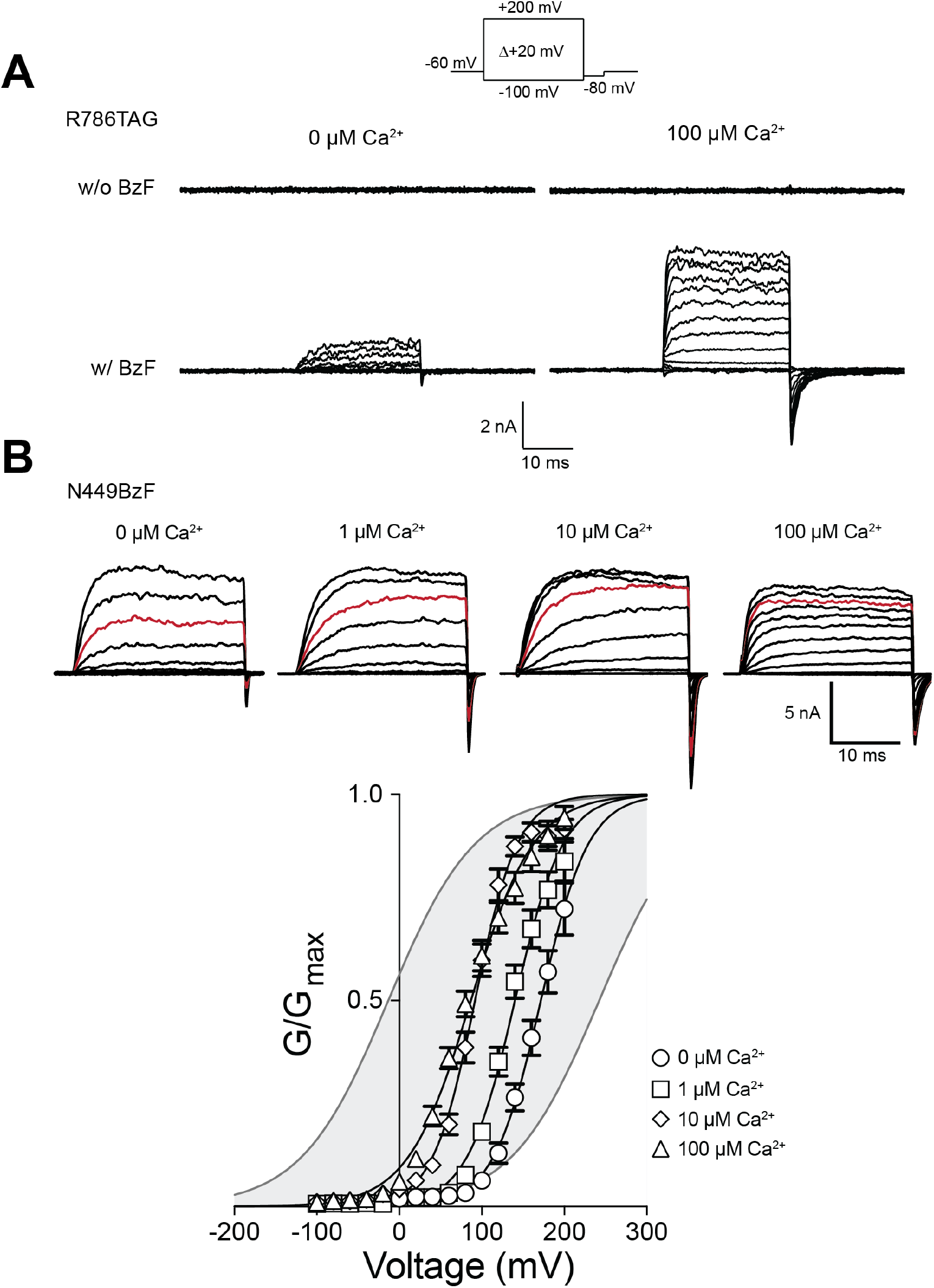
The insertion of BzF in BK-TAG mutants recovers functional and Ca^2+^ sensitive BK channels. (A) The mutant R786TAG showed outward potassium currents in response to a voltage-step protocol only under BzF incubation. This BzF incorporation rendered a channel that retained Ca^2+^ sensitivity as the WT. (B) Incorporation of the bulky BzF residue in some positions altered the Ca^2+^ sensitivity of the mutant channel. In the case of N449BzF, the solely incorporation of this unnatural amino acid produced a compression in the voltage-conductance curves under different Ca^2+^ (top: representative traces of the family of currents under a voltage-step protocol plotted from −100 to + 100. In red is highlighted the +60 mV step for comparison; bottom: summary of the G-V curves under four different Ca^2+^ concentrations for the N449BzF mutant. WT range: shaded gray area).

**Figure S4-1.**
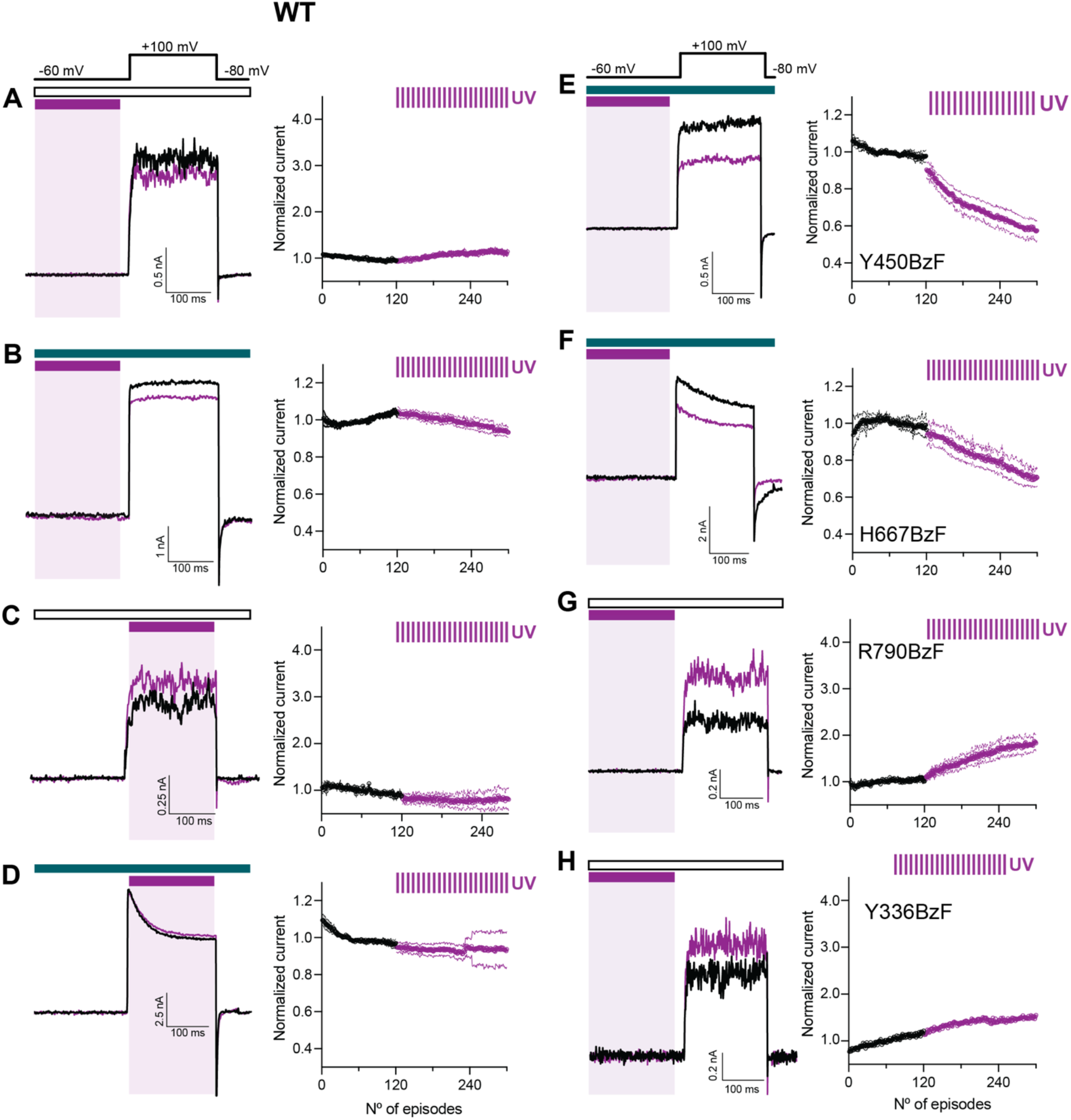
The photocrosslinking effects are specific, and conserved in residues in site A and in site B. (A-D) Subtle to none UV driven effects were observed in the wildtype BK channel (BK667YFP) for all the conditions tested. (E-F) Photocrosslinking under 100 μM Ca^2+^ and hyperpolarized patches only produced a UV-dependent exponential decay in the residue Y450BzF located adjacent to the N449 and not in the H667BzF (notice that the linear decay observed for this last mutant is due to non-specific UV effects or channel rundown, as is similarly observed in the WT). (G-H) Contrarily, photocrosslinking in the site B interface was also observed, although to a lesser extent, in the R790BzF that was not observed in other mutant like Y336BzF, where this BzF-insertion was far from any intersubunit gating ring interface.

**Figure S4-2.**
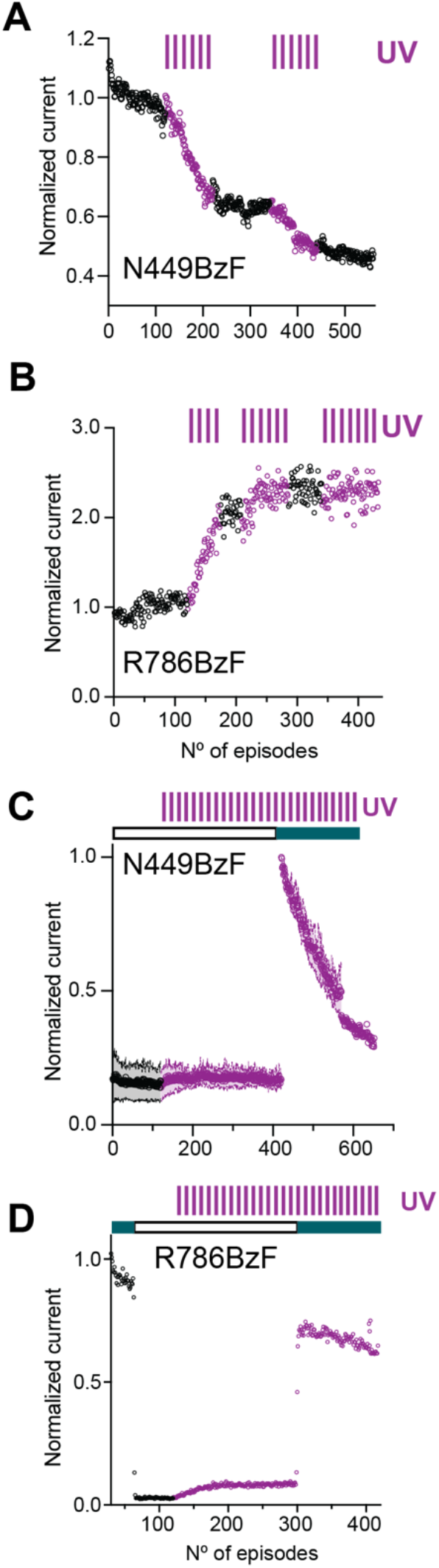
The effects observed for N449BzF and R786BzF are UV and Ca^2+^ specific. (A) Application of alternating periods of UV and darkness showed the current reduction for N449BzF only when the UV was being applied, strengthening the result of a UV specific effect. (B) Similarly to N449BzF, current potentiation of R786BzF was only observed under UV application. (C) N449BzF photocrosslinks only in the presence of 100 μM Ca2+. The application of UV during 0 μM Ca^2+^ did not yield a photocrosslinking not affecting the function of the channel, as the application of 100 μM Ca^2+^ after 0 μM, rendered a similar decay to the one observed in the previous experiments. This indicates that the BzF was not used in the absence of Ca2+. (D) For R786BzF, the application of 0 μM Ca^2+^ is the only one giving the photopotentiation. In this case, it is also important to note that the amplitude of the 100 μM Ca^2+^ showed a rundown but it is linearly related to the baseline at this Ca^2+^ concentration. This remarks the point that the effect R786BzF crosslinking is producing in the BK channel only affects the current of the channels in absence of Ca^2+^.

## Notes

### Competing Interest Statement

The authors have declared no competing interest.

